# HSC-independent definitive hematopoietic cells persist into adult life

**DOI:** 10.1101/2021.12.02.468909

**Authors:** Michihiro Kobayashi, Haichao Wei, Takashi Yamanashi, David J Shih, Nathalia Azevedo Portilho, Samuel Cornelius, Noemi Valiente, Chika Nishida, Wenjin J Zheng, Joonsoo Kang, Jun Seita, Jia Qian Wu, Momoko Yoshimoto

**Affiliations:** Center for Stem Cell and Regenerative Medicine, Brown Institute of Molecular Medicine, University of Texas Health Science Center at Houston, Texas, USA; The Vivian L. Smith Department of Neurosurgery, University of Texas Health Science Center at Houston, Texas, USA; Department of Dept. Biochemistry & Molecular Biology, McGovern Medical School, University of Texas Health Science Center at Houston, Texas, USA; Advanced Data Science Project, RIKEN Information R&D and Strategy Headquarters, Tokyo, Japan; Center for Integrative Medical Sciences, RIKEN, Kanagawa, Japan; Department of Pathology, University of Massachusetts Medical School, Worcester, MA, USA

**Author notes:** Corresponding author, Momoko Yoshimoto MD., PhD.

## Abstract

The stem cell theory that all blood cells are derived from hematopoietic stem cell (HSC) is a central dogma in hematology. However, various types of blood cells are already produced from hemogenic endothelial cells (HECs) before the first HSCs appear at embryonic day (E)11 in the mouse embryo. This early blood cell production from HECs, called HSC-independent hematopoiesis, includes primitive and definitive erythromyeloid progenitors that transiently support fetal blood homeostasis until HSC-derived hematopoiesis is established. Lymphoid potential has traditionally been detected in the extra-embryonic yolk sac (YS) and/or embryos before HSC emergence, but the actual presence of lymphoid progenitors at this stage remains unknown. In addition, whether HSCs in the fetal liver are the main source of innate-like B-1a cells has been controversial. Here, using complementary lineage tracing mouse models, we show that HSC-independent multipotent progenitors (MPPs) and HSC-independent adoptive B-lymphoid progenitors persist into adult life. Furthermore, HSCs minimally contribute to the peritoneal B-1a cell pool; most B-1a cells are originated directly from ECs in the YS and embryo and HSC-independent for life. Our discovery of extensive HSC-independent MPP and B-lymphoid progenitors in adults attests to the complex blood developmental dynamics through embryo to adult that underpin the immune system and challenges the paradigm of HSC theory in hematology.

## Introduction

All blood cells are derived from special endothelial cells (ECs), referred to as hemogenic endothelial cells (HECs) in the extraembryonic yolk sac (YS) and para-aortic region of the mouse embryo during a limited time window^1, 2, 3, 4, 5^. Before the emergence of hematopoietic stem cells (HSCs) from HECs in the aortic regions at E11, multiple waves of blood cell production occur directly from HECs, which contribute to the transient fetal hematopoiesis^4^. During this time, in vitro B-lymphoid potential from HECs in the early YS and embryo has been reported^6, 7, 8, 9^, however, the physiological presence of HSC-independent B-cells has yet to be unequivocally determined. Furthermore, if they exist, how long and to what extent such HSC-independent B-cells persist into postnatal life remains unknown. B-lymphocytes are mainly categorized into three subsets; bone marrow (BM) HSC-derived B-2 cells (e.g., splenic follicular (FO) B-cells), marginal zone (MZ) B-cells, and innate-like B-1 cells that reside mainly in the body cavities. CD5^+^ B-1a cells are not replenished by BM HSCs and have generally been considered to be derived from FL HSCs^10, 11, 12^ whereas contrasting results have also been reported^13, 14^. In addition, it has been reported that progenitors at E10.5 AGM region that have biased B-1 cell potential can acquire B-2 potential upon AGM-derived endothelial niche culture^14^. Therefore, it is plausible that B-2 cells may arise from HECs independently of HSCs.

In this study, using HSC- and EC-lineage tracing mouse models, we found that HSCs in the FL slowly produced MPPs and B-lymphoid progenitors after birth and EC-derived HSC-independent MPPs and B-progenitors persisted in adult for more than 6 months. Furthermore, FL HSCs minimally contributed to the peritoneal B-1a cells and EC-derived HSC-independent B-1a cells were the major population and maintained for life. Transplantation assays of E11.5 HSC-precursors without co-culture demonstrated the presence of transplantable HSC-independent MPPs and B-1a progenitors among this population. Our study resolved the long-lasting controversy of B-1a cells and provide unexpected evidence of HSC-independent MPPs and adaptive B-progenitors in adult life.

## Results

### Fetal HSCs do not contribute to the peritoneal B-1a cell pool in a steady state

*Fgd5* is expressed exclusively in LT-HSCs [lin^-^Sca-1^+^c-kit^+^(LSK)CD150^+^CD48^-^ cells] in the FL and BM. *Fgd5CreERT2:Rosa-TdTomato* (*iFgd5*) mice enable us to label HSCs at a time of Tamoxifen (TAM) injection^15, 16^. We labeled HSCs in E14.5 FL or postnatal day 2 (P2) BM by TAM injection, respectively, and examined Tomato% in various B-cell subsets and BM progenitors over 300 days after birth (Fig. 1A, B, Extended Date Fig. 1A, B). There were variations of Tomato labeling efficiencies among animals and timed matings did not always precisely synchronize the actual embryonic age at the time of TAM administration. Therefore, to evaluate the labeling efficiency in a consistent manner, we calculated a Tomato % ratio of each blood cell type to HSCs (Tomato % ratio =Tomato% of a defined cell type /Tomato% of HSC) as previously described^16^. If a cell population is HSC-derived, Tomato % ratio should become close to 1.0 over time.^16^ Surprisingly, Tomato % ratio of B-1a cells stayed very low (<0.2) up to 300 days after birth (Fig. 1B) even when HSCs were labeled at E14.5 FL stage, indicating that the majority of adult peritoneal B-1a cells were HSC-independent. Furthermore, the Tomato % ratios of MPPs and FO B-cells showed 0.7-0.8 and 0.5, respectively (Fig. 1B, Extended Date Fig. 1A, B). This result raised a question as to whether some of these cells in adults arise independently of HSCs.

**Figure 1.**
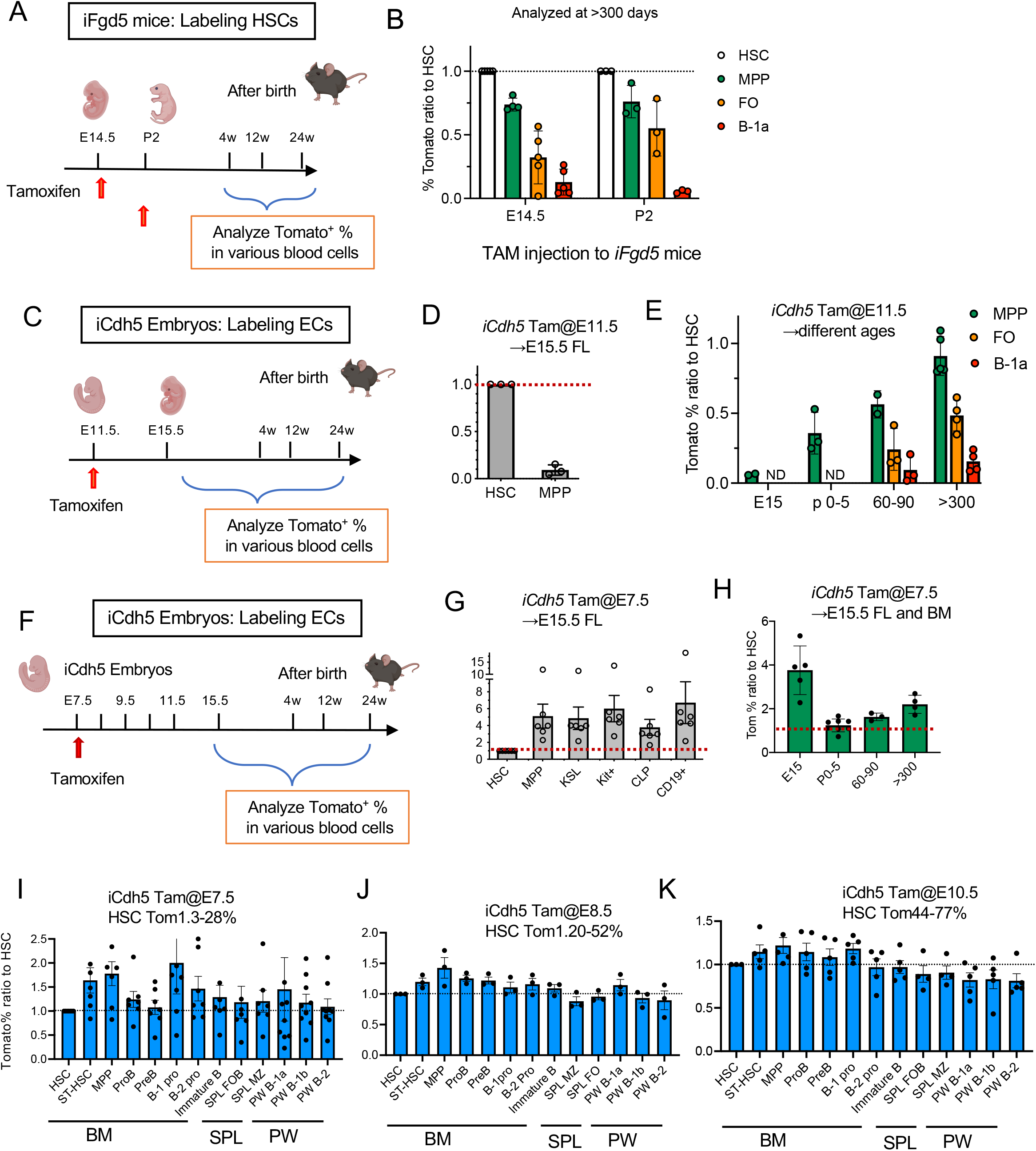
B-1a, MPP, and other lymphoid cells arise independently of HSCs and persist into adult. (A) TAM was injected once at E14.5 or P2 into *iFgd5* mice to label HSCs. Tomato^+^ blood cells in the BM, spleen, and peritoneal cavity were examined at different time points after birth. (B) The relative Tomato% ratios of MPP, Follicular (FO) B cells, and B-1a cells to HSCs are shown. TAM was injected to *iFgd5* mice at E14.5 or P2 and mice were examined more than 300 days after birth (n=3-5). (C) TAM was injected into *iCdh5* pregnant mice at E11.5. Tomato^+^ HSPCs were examined at different time points such as E15.5 and after birth. (D) The relative Tomato % ratio of MPPs to LT-HSCs in E15.5 FL when ECs were labeled at E11.5 (n=4). (E) The relative Tomato% ratios of MPPs, splenic FO, and peritoneal B-1a cells to LT-HSCs at different time points (P0-5, days 60-90, days 300<, n=3-6 at each time point). ND: not done. (F) TAM was injected into *iCdh5* pregnant mice at E7.5 and Tomato^+^ HSPCs were examined at different time points such as E15.5 and after birth. (G) The relative Tomato% ratio of each target cell population to LT-HSCs in E15.5 FL, when ECs were labeled at E7.5 (n=6). (H) The relative Tomato% ratio of MPPs to LT-HSCs in the FL and post-natal BM at different time points (P0-5, days 60-90, days 300<, n=3-5). The relative Tomato% ratios of HSPCs and B cell subsets to LT-HSCs at >300 days after birth when TAM was injected at E7.5 (I), E8.5 (J), and E10.5 (K). N=3-7 for each TAM injection. BM: bone marrow, SPL: spleen, PW: peritoneal wash.

### MPPs and B-progenitors in the FL are HSC-independent and originated at as early as E7.5

It has been reported that FL MPPs, but not LT-HSCs, produced B-1a cells most efficiently upon transplantation^13, 14^. Since fetal HSCs were not the major drivers of the peritoneal B-1a cells in steady states (Fig. 1B), FL MPPs that have B-1a potential must be derived from precursors at earlier stages than FL HSCs, such as HECs that can produce various hematopoietic cells^17, 18^. Therefore, we sought the origin of FL MPPs by using an EC-lineage tracing mouse model. *Cdh5* is a specific marker of ECs and *Cdh5CreERT2: Rosa-TdTomato* mice (*iCdh5*) mice are widely used to label ECs at a time of TAM injection^19^. First, we tried to label HECs that produce the first de novo HSCs at E11.5. EC-labeling at E11.5 exclusively marked HSCs when we analyzed E15.5 FL (Fig. 1C, D). While 18.8±12.7% of LT-HSCs were Tomato^+^, only 2.3±2.8% of MPPs were Tomato^+^, and its Tomato% ratio was only 0.1 (n=3) (Fig. 1D). These results indicate that even the first HSCs produced at E11.5 do not yet differentiate into MPPs in the E15.5 FL, but gradually produce MPPs and FO B-cells after birth (Fig. 1E). Additionally, B-1a cells were not efficiently labeled by E11.5 TAM injection even when analyzed at >300 days (Fig. 1E, Extended Date Fig. 1C), in line with the HSC-lineage tracing results that HSCs do not produce B-1a cell efficiently.

Next, we sought the origin of FL HSC-independent MPPs at the earlier embryonic stages before HSC emergence. We labeled ECs at E7.5 and examined Tomato^+^ MPPs and other hematopoietic progenitors in E15.5 FL (Fig. 1F). Surprisingly, whereas LT-HSCs were barely labeled, around 10-30% of MPPs were Tomato^+^ (Fig. 1G, Extended Date Fig. 1D). Furthermore, other hematopoietic progenitors including common lymphoid progenitors (CLPs) and CD19^+^ B-progenitors showed higher Tomato% than that of LT-HSCs (Fig. 1G, Extended Data Fig. 1D). Because we labeled ECs, not HSCs, in *iCdh5* mice, the Tomato ratio=1.0 indicates that the target cells and HSCs are derived from ECs at the same stage, and >>1.0 or <<1.0 indicates that these two populations are derived from ECs at different time points^20^. The Tomato % ratios of MPP, CLP, and B progenitors to HSCs were much >> 1.0 (Fig. 1G, H), indicating that E15.5 FL MPPs and B-lymphoid progenitors were HSC-independent. Importantly, these HSC-independent MPPs and B-progenitors marked at E7.5 persisted into adult life, more than 300 days after birth (Fig. 1H, I).

When E9.5 ECs were labeled, the Tomato % of MPPs, other progenitors, and HSCs in the E15.5 FL were similar (the ratio was near 1.0) (Extended Data Fig.1E). Considering that HSCs expand rather than differentiate during E10.5 to E15.5^21, 22^, this result suggests that most progenitors and HSCs were simultaneously produced from HECs at E9.5. Importantly, these E7.5 and 9.5 HSC-independent MPPs persisted more than 300 days after birth because their tomato ratio kept >1.0, suggesting that HSC-independent MPP-derived hematopoiesis occurs even in the adult BM (Fig. 1H, I, Extended Data Fig. 1F).

### FL MPPs contain HSC-independent common B-1 and B-2 progenitors

As Tomato^+^E15.5 FL MPPs are HSC-independent (Fig. 1G) and contain B-1a precursors upon transplantation^14^, we examined their B-1 and B-2 progenitor potential using modified B-cell colony assays^14^. From 500 Tom^+^ MPPs marked at E7.5 in *iCdh5* mice, 29 B-progenitor colonies were detected (Extended Data Fig. 1G). Among them, we found 20 B-1 progenitor colonies and 9 B-1 and B-2 progenitor mixed colonies. These data showed that E15.5 FL HSC-independent MPPs contain B-1 progenitors and common lymphoid progenitors that can differentiate into both B-1 and B-2 cells.

### Most B-1a cells are derived from E7.5-10.5 HECs

Since HSCs after E11.5 showed minimal contribution to the peritoneal B-1a cells (Fig. 1E), we examined at which stage of HECs mark the peritoneal B-1a cells most efficiently. We expected that B-1a cells were derived from ECs at early embryonic stages similar to brain macrophage (Extended Data Fig. 1H)^20^. However, B-1a cells were marked by ECs during E7.5 to E10.5 (Extended Data Fig. 2, 3). Other B-cell subsets including FO and MZ B cells showed similar labeling patterns (Extended Data Fig. 2, 3). Tomato% ratios of these B-1a and B-2 cells to HSCs were just slightly higher than 1.0 when ECs were labeled at E7.5 or E8.5 (Fig. 1I, J). When E10.5 ECs were labeled, almost all B-lymphoid subsets including B-1a cells showed similar Tom % with that of HSCs (the ratio of nearly 1.0) (Fig. 1K). These results suggest that B-1a cells, a part of other B-cell subsets, and HSCs were produced simultaneously from ECs because HSCs after E11.5 contributed to only a part of each B-cell subset (Fig. 1E, Extended Data Fig. 1C). Taken together, the data indicate that most B-1a cells are derived from ECs during E7.5 to 10.5, and a portion of FO and Marginal zone B-cells are also HSC-independent.

In order to explain the results from the *iCdh5* mouse (Fig. 1I-K, Extended Data Fig. 1C), we constructed a mathematical model of label tracing experiments by extending a previously established model^23^. We constructed three variants of the label tracing model based on competing hypotheses for the cell differentiation tree (Extended Data Fig. 4A). The base model *M_0_* assumes a linear differentiation path from hemogenic EC via HSC, MPP, B-1 progenitor, and finally to B-1 cell. The *M_1_* model hypothesizes that ECs can directly differentiate into MPPs. In addition to *M_1_* model, the *M_2_* model hypothesizes that ECs can directly differentiate into B-1 progenitors. After model fitting, the *M_2_* model yielded label tracing predictions that most closely resemble the experimental label tracing data from the *iCdh5* mouse (Fig. 1I-K, Extended Data Fig. 1C, 4, 5). These results indicate that the differentiation tree of the *M_2_* model can best explain the experimental data, providing additional support that ECs directly differentiate into MMPs and B-1 progenitors independently of HSCs during fetal development.

### Single cell-RNA-sequencing showed heterogeneity and B-lymphoid signatures of pre-HSC and HSC population

Lineage tracing studies demonstrated the multiple waves of HSC-independent hematopoiesis including MPPs, B-progenitors, and B-1a cells. We recently reported that E10.5 HSC-precursor (pre-HSC) population shows B-1 biased repopulating ability^14^. Pre-HSCs are intermediate precursors between HECs and adult repopulating HSCs detected during E10.5 to 11.5 and express VE-cadherin (VC), encoded by *Cdh5*^24^. At E11.5, the first adult repopulating HSCs are detected^24^. To understand the transition from pre-HSCs to HSCs or MPP, and their heterogeneity of hematopoietic capability, we performed single-cell (sc) RNA-sequencing of E11.5 AGM and YS VC^+^c-kit^+^EPCR^+^ pre-HSC population, E12.5 FL HSC (CD45^+^LSK^+^EPCR^+^) and E14.5 FL HSCs (CD45^+^LSK^+^CD150^+^CD48^-^) (Extended Data Fig. 6). In parallel, selected sorted subsets were transplanted into sublethally irradiated NSG neonates to validate their hematopoietic capabilities (Extended Data Fig. 6D).

We sorted individual cells from above populations and generated single-cell full length transcriptome using SMART-seq. We distributed the genes (read counts >10) in each cell and excluded the cells which expressed less than 2000 genes and genes that were detected in less than 10 cells (Extended Data Fig. 6E). At last, 95 cells and 11,814 genes were used for further analysis. PCA analysis of scRNA-seq showed clear separation of FL HSCs from pre-HSCs (Fig. 2A). Notch-related genes were all highly expressed in many AGM cells but were downregulated in FL HSCs (Extended Data Fig. 7) as previously reported transitional requirement of Notch signaling^25^. Unbiased sc consensus clustering (SC3) of the whole transcriptomes separated these cells into four distinct clusters (Fig.2B). The top 10 differentially expressed genes are depicted in Extended Data Fig. 8. We focused the expressions of HSC- and lymphoid cell-related genes in each cell (Fig. 2C) and examined the trajectory of cell states with ordering of cells from E11.5 AGM to E14.5 FL HSCs (Extended Data Fig. 9) using pseudo-time analysis. Trajectory map indicated the progression of E11.5 AGM pre-HSCs to E12.5 and E14.5 FL HSCs (Extended Data Fig. 9). Almost all pre-HSCs and HSCs expressed essential genes for B-cell development, such as *Ikz1* and *Tcf3* in addition to many HSC-related genes (Fig. 2C, Extended Data Fig. 9). *Bcl11a*, important for fetal and adult B cell development and globulin switching, is also widely expressed among pre-HSCs and HSCs (Fig. 2C, Extended Data Fig. 9). Interestingly, essential BCR signaling related genes, such as *CD79b and Btk*, were heterogeneously expressed in all cell types, even in FL HSCs, suggesting their biased B-lymphoid potential (Fig. 2C, Extended Data Fig. 9). *Lin28b*, encoding RNA-binding protein and critically important for B-1a cell generation^11, 26^, is also heterogeneously expressed among pre-HSC and HSC populations (Extended Data Fig. 9), which may explain the contrasting results of FL HSCs transplantation assays^11, 12, 13, 14, 27^. *Pbx1*, the Hox cofactor and proto-oncogene, is essential for B-lymphoid lineage commitment and HSC-self-renewal/maintenance^28, 29, 30^. While *Pbx1* was widely expressed among pre-HSCs, its expression was only seen in a small portion of HSCs (Fig. 2C, Extended Data Fig. 9). These data displayed the heterogeneity of highly purified pre-HSC and HSC populations and even genes that have been considered critical for HSC maintenance may not be expressed ubiquitously in FL HSCs. We also performed the velocity analysis^31, 32^ to understand the gene expression status of each cell. Interestingly, the velocity analysis showed two directions of the gene expression status into the left and the right shown in Fig. 2D. When we compared the gene expressions between the cells on the left and right sides, 41 genes were differentially expressed (supplementary file). We examined these gene expressions in BM HSPC and B-progenitors from the database at ImmGen (Fig. 2E) and found the difference of these gene expressions reflected the ones found in BM B-progenitors or HSPCs. This result suggests that there may be divergence into HSC and B-cell commitment of pre-HSC and HSC populations.

**Figure 2.**
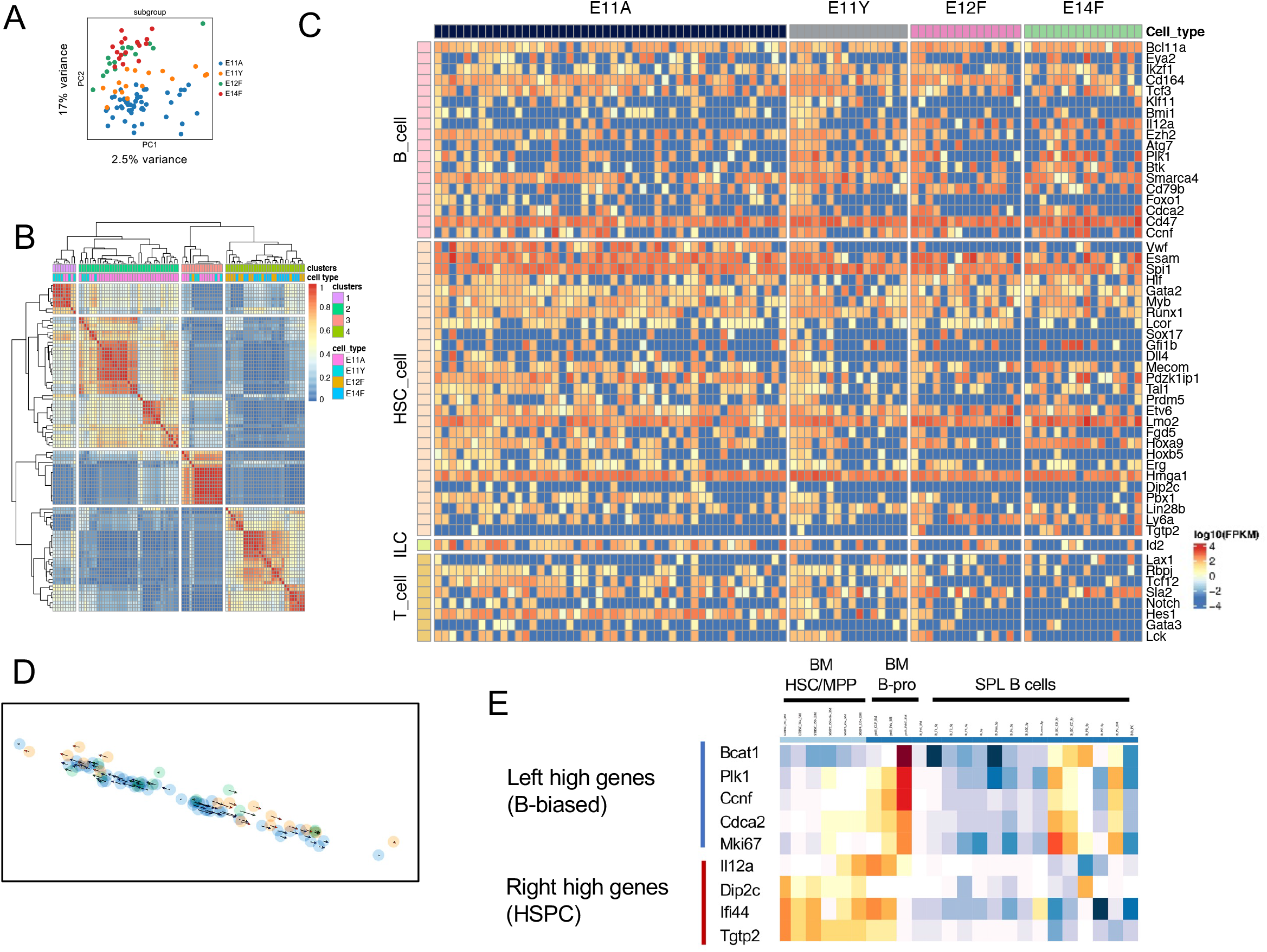
scRNA-seq analysis showed HSC and B-lymphoid signatures in of E11.5 pre-HSCs and E12&14 FL HSCs. (A) Dimensionality reduction of scRNA-seq data using PCA colored by cell type. E11A, E11.5 AGM pre-HSC, E11Y, E11.5 YS pre-HSC, E12F, E12.5 FL HSC, E14F, E14.5 FL HSCs. (B) SC3 consensus matrix predicted 4 clusters. (C) A heat map depicting the expression of HSC, B, T, and ILC related genes in E11 AGM&YS pre-HSC populations and E12.5&14.5 FL HSCs. The red, blue, and yellow intensities indicate high, low, and intermediate expression levels, respectively. (D) Velocity analysis of scRNA-seq of E11.5 pre-HSC and E12.5 &14.5 HSCs. Small arrows show the direction of the velocity of single cells. (E) Heat map of gene expressions detected in right and left directions in the velocity analysis, which were applied to gene expressions in the BM HSPC and B-progenitors using the database at ImmGen (https://www.immgen.org/).

### HSC-independent MPPs and B-1 biased repopulating cells are present in E11.5 Embryo

Upon the heterogenous gene expression signatures of HSPC and B-cell lineages in pre-HSC population, we examined the presence of HSC-independent MPPs and B-1a precursors in the pre-HSC population in transplantation settings. We injected 5 −10 cells of CD45^+^Ter119^-^VC^+^c-kit^+^EPCR^+^ pre-HSC population isolated from E11.5 aorta-gonad-mesonephros (AGM) region (Extended Data Fig. 6A) into sublethally irradiated NSG neonates. Of total 25 recipient mice, 15 showed donor-derived CD45.2^+^ cells (>0.1%) in the peripheral blood (PB) (Fig. 3A). When we analyzed the transplanted mice at 4-6 months post transplantation (Fig. 3B-D), we found there were four types of engraftment patterns; multi-lineage engraftment including BM LSK cells (HSC-engraftment, Fig. 3B-D, E-G, mouse #1-5); multi-lineage engraftment without BM LSK cells (MPP engraftment, Fig. 3B-D, H, I, J mouse #6-11); and only B-1 and B-2 cell engraftment (Fig. 3B, C, mouse #12); and only B-1 cell engraftment (Fig. 3B, C, K, mouse #13-15).

**Figure 3.**
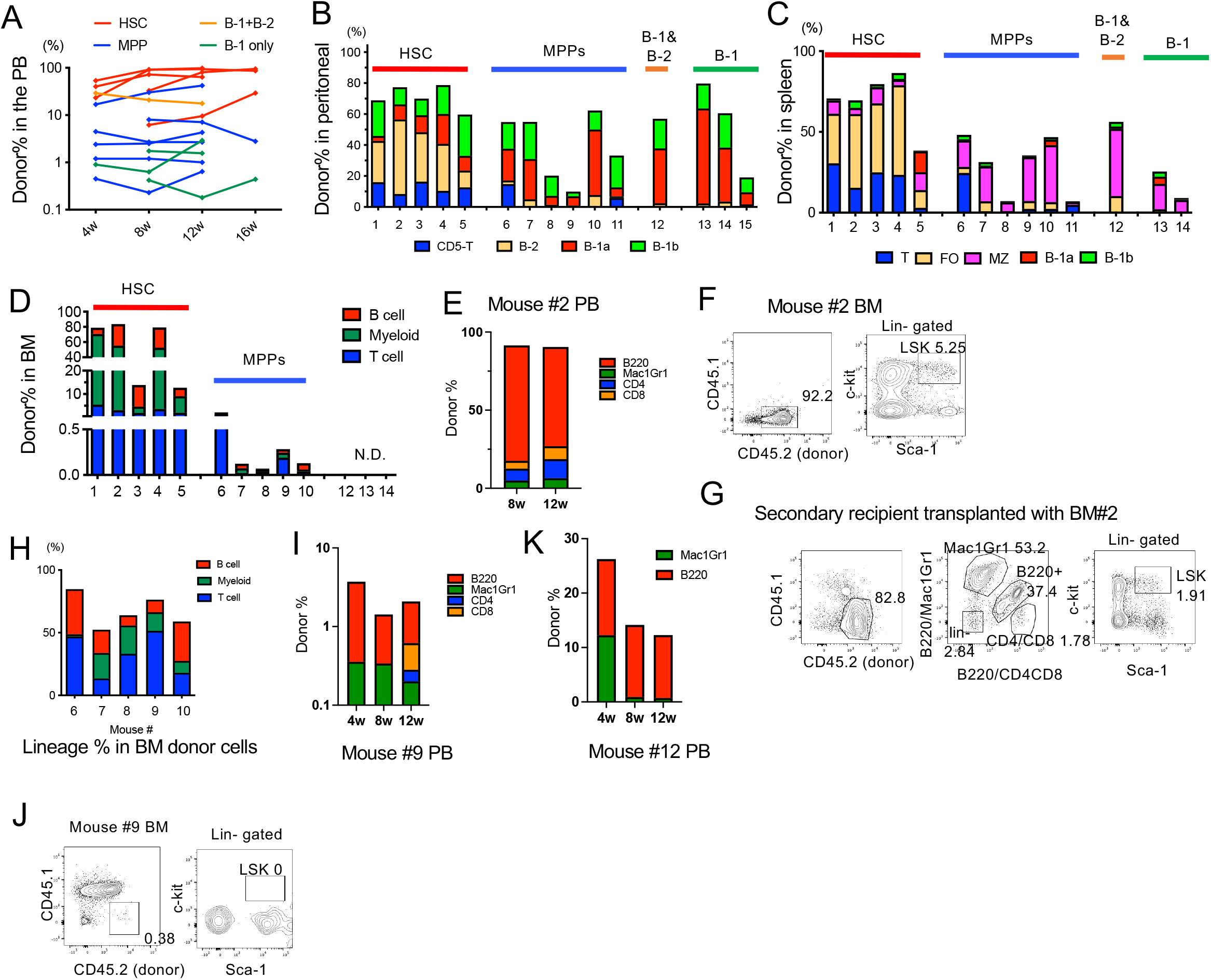
LT-HSCs, MPPs, and B-1 repopulating cells arise independently from CD144^+^c-kit^+^EPCR^+^ cells in E11.5 AGM region. Five to fifty pre-HSCs from E11.5 AGM region were injected into sublethally irradiated NSG neonates. (A) CD45.2^+^ donor cell % in the peripheral blood of the recipient mice 4-6 weeks after transplantation. Donor cells % and their composition within the lymphoid subsets in the peritoneal cells (B), spleen (C), and BM (D) are depicted. The donor-derived cell lineages in the recipient PB during the time course of mouse #2 (E, HSC-engrafted), mouse #9 (I, MPP-engrafted), and #12 (K, B-1 and B-2 cell-engrafted). (H) The % of donor-derived lineages in the recipient BM of mouse #6-10 are depicted. Although the donor cell % was low, multi-lineage repopulation was observed in the recipient BM. The representative FACS plots for donor LSK cells in the first (F, mouse #2 and J, mouse #9) and secondary recipient BM (G).

Mice #1-5 showed long-term multi-lineage repopulation with significant donor cell % in the PB, peritoneal cavity, spleen, and BM (Fig. 3A-E) and predominant B-2 cell engraftment with B-1a and B-1b cells (Fig. 3B). In the recipient BM, successful donor derived LSK cell repopulation was also confirmed (Fig. 3F). Thus, these mice were categorized as HSC-repopulated mice. Importantly, the secondary recipient BM also showed donor-derived LSK repopulation (Fig. 3G), indicating that LSK-repopulating cells are the functional HSCs that harbor a self-renewal ability. In contrast, mouse #6-11 showed long-term reconstitution without donor-LSK cells in the BM (Fig. 3A-D, H, I, J), indicating that they were engrafted with HSC-independent progenitors, thus named MPP-engrafted mice. These mice showed predominant B-1 and MZ B cell engraftment with seemingly diminishing B-2 cells in the peritoneal cavity and spleen (Fig. 3B, C) and B-2, T, and myeloid cell engraftment in the PB and BM (Fig. 3D, H, I, J), although the donor percentage in the BM was very low (<0.5%) (Fig. 3D, J). Mouse #13-15 showed B-1a, B-1b, and MZ B-cell engraftment in the peritoneal cavity and spleen but did not show donor-derived cells in the BM (Fig. 3B-D), indicating HSC-independent B-1 cell engraftment. One mouse (#12) showed only B-1 and B-2 cell engraftment (Fig. 3B-D, K), suggesting the presence of common B-1 and B-2 progenitors.

These results strongly indicate that HSC-independent long-term engraftable MPPs and B-1 precursors are present in E11.5 AGM region, also in line with the scRNA-seq data showing the heterogeneity of these cells. Importantly, all engrafted mice showed B-1a cell repopulation whereas B-2 cells were dominant in HSC-engrafted mice; thus, it seems that B-1a potential is the default within E11.5 pre-HSCs and B-2 cell dominant capacity is a hallmark of functional HSCs.

## Discussion

We demonstrated that definitive hematopoietic cells including MPPs, all B-cell subsets, and HSCs are independently produced from HECs and persist into adult life. While HSC-independent B-1a cells are rarely replaced by HSC-derived cells, HSC-independent MPPs and B-2 cells are gradually replaced by HSC-derived cells. These findings challenge the current paradigm of HSC-derived hematopoiesis and finally clarified the longstanding unresolved question regarding the origin and main source of B-1a cells.

In adult murine hematopoiesis, it has recently been reported that MPP is a main driver of native hematopoiesis and the discrepancy of HSC- and MPP-derived clones have been observed^33^. Our results explain this discrepancy, where the majority of MPPs in young mice are HSC-independent, derived from fetal ECs, and HSC-derived hematopoiesis appears later, This is also in line with the previous report that postnatal HSC-labeling showed gradual increase of HSC-derived-lymphocytes over 32 weeks^16, 23, 34, 35^ and a recent report showing minimal contribution of definitive HSCs to fetal hematopoiesis in a fish model^36^. In addition, the presence of HSC-independent MPPs in the pre-HSC population has also been reported using AGM-EC co-culture system^18^. Therefore, together with our results, pre-HSC population is essential not only for maturing to HSCs but also containing MPPs that support hematopoiesis in postnatal life.

Our lineage tracing results (Fig. 4A), difficult to reconcile with the current classic model (Fig. 4B), instead lead us to propose a multiple hematopoietic wave model (Fig. 4C). In the current classical model (Fig. 4B), YS-derived EMPs maintain hematopoietic homeostasis until perinatal periods^37^. Once HSCs are produced in the embryo, HSCs expand in the FL, start HSC-derived hematopoiesis, and migrate to the BM just before birth, where HSCs maintain hematopoiesis for life. Our model proposes that multiple waves of HSC-independent hematopoiesis, including EMP, B-1 precursor and MPP production, occur from ECs, and persist into adult life. HSCs are also produced as the final wave of EC-derived blood production. While HSCs expand in the FL, HSC-derived hematopoiesis seems to first start after settling the BM, and gradually replaces the HSC-independent hematopoietic cells over time. B-1a precursors are produced directly from ECs or via HSC-independent MPPs at early- and mid-gestation but not replaced by HSC-derived cells in steady state.

**Figure 4.**
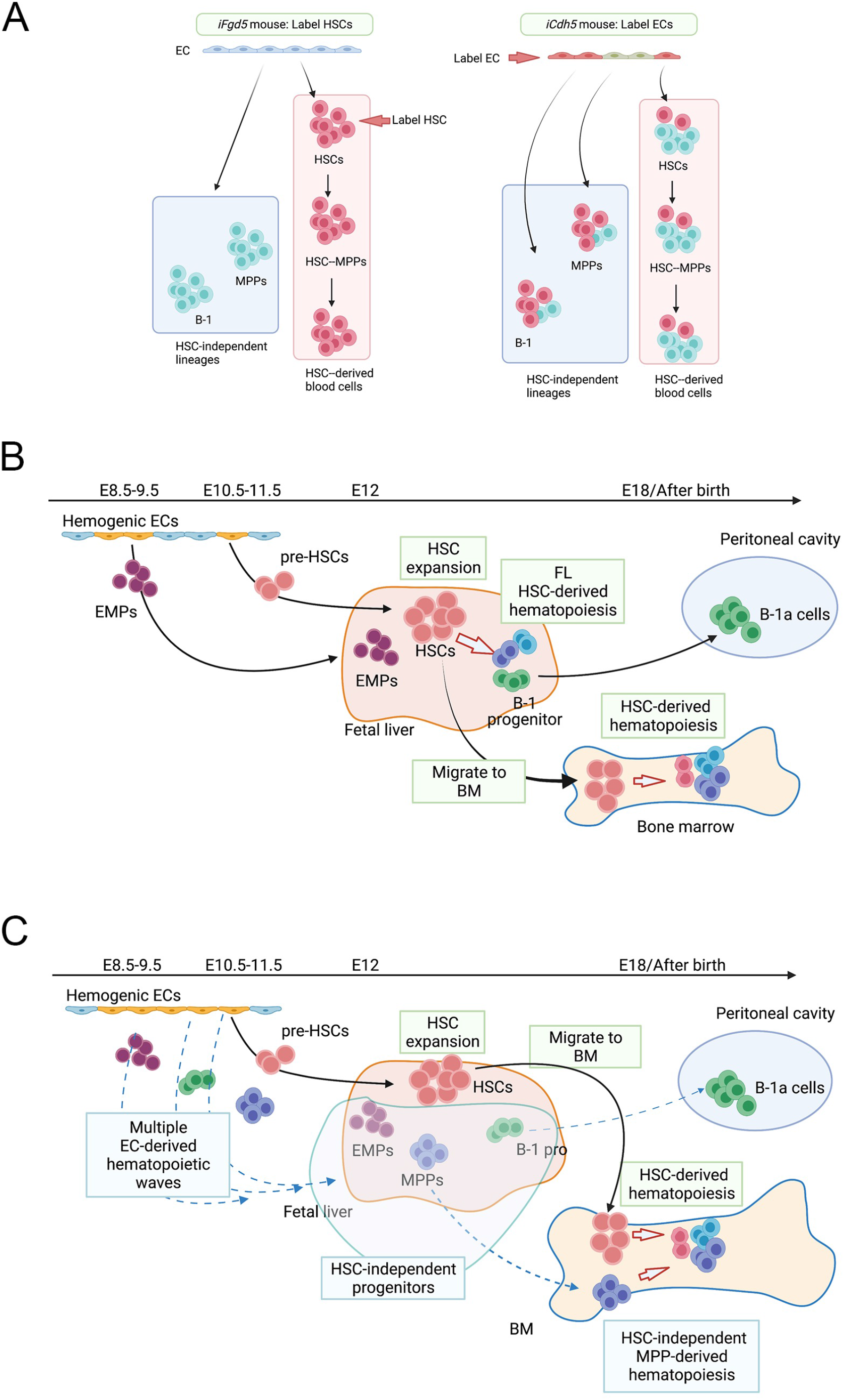
The current and proposed models for developmental hematopoiesis in the mouse embryo based on the lineage tracing studies. (A) The summary of the results from HSC- and EC-lineage tracing studies. While HSC-lineage tracing does not label HSC-independent blood lineages (left), EC-lineage tracing labels blood cells with different percentages depending on the timing when those blood cells are produced from ECs (right). EC-labeling at E7.5 marked more HSC-independent blood cell types than HSCs. (B) The current classical hematopoiesis model during fetus. Hemogenic ECs produce EMP, possibly lymphoid progenitors (not indicated), and HSCs. These EMPs and HSCs seed the fetal liver where EMPs provide mature definitive erythroid and myeloid cells and HSC-derived hematopoiesis start while HSC self-renew and expand at the same time. (C) Our working model proposing EC-derived multiple waves of fetal hematopoiesis. Almost all hematopoietic progenitors including EMP, MPP, and B-1 progenitors are produced from hemogenic ECs during E7.5-10.5 independently of HSCs. HSC production is the final wave of EC-derived hematopoiesis. These progenitors and HSCs seed the fetal liver and then bone marrow before birth. While HSCs mainly self-renew and expand in the fetal liver, EC-derived (HSC-independent) blood progenitors maintain hematological homeostasis and provide mature blood cell subsets until HSC-derived progenitors replace them in postnatal life. However, peritoneal B-1a cells are not replaced by HSC-derived progenitors and maintain themselves for life.

There are some discrepancies between lineage tracing study and transplantation assays regarding the B-1a cell potential. While E11.5 EC labeling did not mark B-1a cells, E11.5 VC^+^ pre-HSCs showed B-1a repopulating ability. This may be due to the difference of expression timings of Cdh5 transcriptome and VC surface protein. In addition, the first HSCs at E11.5 possess B-1a repopulating ability upon transplantation whereas FL HSCs did not. Therefore, HSCs may have B-1a cell potential within a limited time window in transplantation or stress settings.

Taken together, we unveiled unappreciated presence of HSC-independent hematopoietic progenitors in adult life and challenge the paradigm of HSC-dogma in hematology.

## Acknowledgement

The scRNA-sequencing work was performed at the Single Cell Genomics Core at BCM partially supported by NIH shared instrument grants (S10OD023469, S10OD025240) and P30EY002520.This work is supported by NIH R01AI121197 (M.Y.), R01AI147685 (J. K.). H. W. and J. Q. W. are supported by grants from the National Institutes of Health R01 NS088353 and R21 NS113068-01. This work was also partly supported by the National Institutes of Health (NIH) through grants 1UL1TR003167 and the Cancer Prevention and Research Institute of Texas through grant RP170668 (W.J.Z).

Some figures were created with BioRender.com.

## Data availability

Sequencing data have been deposited in the GEO database under accession number GSE182206.

All original code has been deposited at github and is publicly available as of the date of publication. URLs are listed in the key resources table.

## Author contributions

M.K. conceived, design, and performed experiments and analyzed the results. H.W., T.Y., J.S. and J.Q.W participated in bioinformatics data analysis. D.J.S and W.J.Z calculated the mathematical model, N.A.P., S.C., N.V., C.N. performed experiments, J.K. analyzed the results and edited the manuscript, M.Y. conceived, design, and performed experiments, analyzed the results, wrote and edited the manuscript.

## Methods

### Experimental Animals

Cdh5(PAC)-CreERT2 mice were obtained from Dr. Ralf Adams. Fgd5CreERT2 mice (Stock No: 027789) and Rosa-TdTomato mice (Stock No: 007909) were obtained from Jackson Laboratory. Cdh5(PAC)-CreERT2 mice were crossed with Rosa-TdTomato mice and Cdh5CreE2:Rosa-Tomato mice were generated. Similarly, Fgd5CreERT2 mice were crossed with Rosa-TdTomato mice and Fgd5CreERT2:Rosa-Tomato mice were generated. These mice were timed mated with Rosa-Tomato mice and the vaginal plugs were confirmed in the following morning. The noon on the day that the plug was found was counted as embryonic day 0.5. Tamoxifen (15ng/mother body weight) was administrated into timed mated pregnant iCdh5 dams at E7.5, 8.5, 9.5, 10.5, 11.5 respectively, or into timed mated pregnant Fgd5 dams at E14.5, or P2 neonates. 10-15 mice for each Tam injection date were examined.

For transplantation assays, C57BL/6 mice were timed mated and embryos at E11.5 were harvested from the pregnant dams. AGM region was dissected from the embryo and lin-CD144+c-kit+EPCR+ cells were sorted for donor cells. The embryonic age was confirmed by the somite numbers and developmental features of the embryos.

NOD/SCID/Il2γc^null^ mice (NSG mice. Jackson Laboratory Stock No: 005557, OD.Cg-*Prkdc^scid^ Il2rg^tm1Wjl^*/SzJ mice) were timed mated and day2-5 neonates were used for recipients of transplantation assays. Recipient NSG neonates were sublethally irradiated (150rad) before donor cells were injected into facial vein. For secondary transplantation, 1-2 million BM cells from the primary recipient mice were injected into lethally irradiated adult BoyJ mice.

Mice were kept in specific pathogen free condition and all the experimental procedures using the mice were approved by Animal Welfare Committee at UTHealth.

### Lineage tracing experiments

#### Cdh5CreERT2

Tosa-Tomato mice were timed mated. A single dose of Tamoxifen (TAM)(Sigma) 15ng/mother body weight together with Progesterone (7.5ng/mother body weight) solved in corn oil was administrated to the pregnant dam by oral gavage at E7.5, 8.5, 9.5, 10.5, and 11.5 respectively. TAM usually makes the delivery difficult, therefore, Cesarean section was performed on day 19 pregnant dams to rescue the embryos and these pups were taken care of a surrogate mother prepared in advance. Fgd5CreERT2: Rosa-Tomato mice were used for marking HSCs, therefore, TAM was injected into E14.5 pregnant dam or P2 neonatal mice. We harvested various hematopoietic tissues including peritoneal cells, spleen, thymus, and bone marrow from TAM administrated embryos/mice and examined Tomato^+^ percentages in each hematopoietic subset. We compared relative Tomato+ percentage between the cell population of interest and HSCs. The surface markers used for flow cytometry is listed in the table.

### Mathematical modeling of label tracing data

We extended a previously established mathematical model for label tracing data^23^. Consider three successive cell compartments in hematopoiesis: upstream *u*, reference *r*, and downstream *d*. For example, in the differentiation path HEC → HSC → MPP, compartment *u* represents hemogenic EC, compartment *r* represents HSC, and compartment *u* represent MPP. The change in cell number *N_r_* over time *t* is given by:

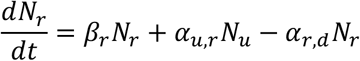

where *β_r_* is the net proliferation rate, *α_u,r_* is the differentiation rate from compartment *u* to *r*, and similarly for *α_r,d_*. Solving this differentiation equation leads to an exponential growth model for *N_r_* (when differentiation rates are set to zero). Biologically, over the lifetime of an organism, a logistic growth model may be more realistic because each cell type would have an upper limit on population size. We therefore modify the label tracing model as follows:

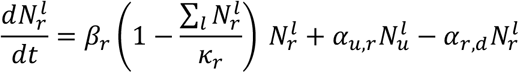

which imposes a carrying capacity *κ_r_* that modulates the net proliferation rate. Integer *l* indexes the cell number *N_r_* in order to distinguish label compartments (Tomato^-^ vs. Tomato^+^). The label compartments share the same parameters (*β_r_, α_u,r_, α_r,d_*), because these parameters depend on the cell type and are independent of label status. Conversely, both label compartments simultaneously experience a common population limit, because the proliferation rate is determined by the total cell number 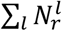 marginalized over label status, as shown in the above equation.

We implemented discrete-time simulations for label tracing models with various differentiation trees (*M*_0_, *M*_1_, *M*_2_) in the R environment (v4.1.1). The *κ* parameters were initialized to steady-state compartment sizes that were previously determined whenever available (Busch et al., 2015). The simulation models were used to predict cell numbers across time for each cell type and label compartment, from which the label proportions and label ratios were calculated. The model parameters were tuned so that the predicted label ratios resemble the experimentally determined Td-Tomato label ratios.

### B-progenitor colony forming assay

Five hundred FL MPPs were plated to methylcellulose containing 10ng/ml IL-7 with 10^5^ OP-9 cells. Eight days after plating, colony numbers were counted and each colony was picked up and stained with anti-mouse CD45, AA4.1, CD19, B220, and CD11b to identify B-1 and B-2 progenitors using flow cytometry.

### scRNA-sequencing

E11.5 pre-HSCs, E12.5 and 14.5 FL HSCs were single cell sorted into 96 well plate (1 cell /well). RNA were extracted from each well and converted into cDNA using SAMRT-Seq Single cell kit (Takara). DNA library was made and sequenced at Single Cell Genomic Core at the Baylor College of Medicine. Briefly, following cDNA synthesis, Nextera XT DNA library preparation kit (Illumina) was used to prepare library.120pg of cDNA was simultaneously fragmented and tagged with adapter sequences by transposome. The product was then amplified using 12 cycles of PCR and purified. Final library was sequenced using Illumina Novaseq 600.

### Bioinformatics analysis of scRNA-sequencing

The quality of all sequenced samples was analyzed using FastQC. Raw reads were aligned to the GRCm38 reference genome using STAR with default parameters. The expression count matrix was generated using htseq-count. We filtered genes whose read counts less than 10 and cells that were less than 2000 genes. Read counts were normalized using DEseq2 with default parameters. The normalized matrix was clustered by SC3. We chose K=4 for SC3 as the best represented the heterogeneity in our dataset. Marker genes in each cluster were filtered by the area under the ROC curve (auroc) > 0.85 and the adjusted p-values<0.01. Trajectory and pseudotime analysis were performed by monocle2 package with default parameters.

PCA analysis was performed using scikit-learn software package (https://scikit-learn.org/stable/). RNA Velocity analysis was performed according to the original article^31, 32^. Unspliced pre-mRNA counts and mature spliced mRNA counts for each cell were computed from the BAM files generated above., then RNA velocity was computed using scVelo software with default parameters. The result was visualized onto the PCA plot based on the expression profile of mature spliced mRNA for each cell. Each arrow represents a direction of a cell transition based on the RNA Velocity.

### Statistical analysis

Non-parametric student-t test was used for statistical analysis.

