## Supplementary Fig. 1- 9 for "HSC-independent definitive hematopoietic cells persist into adult life"

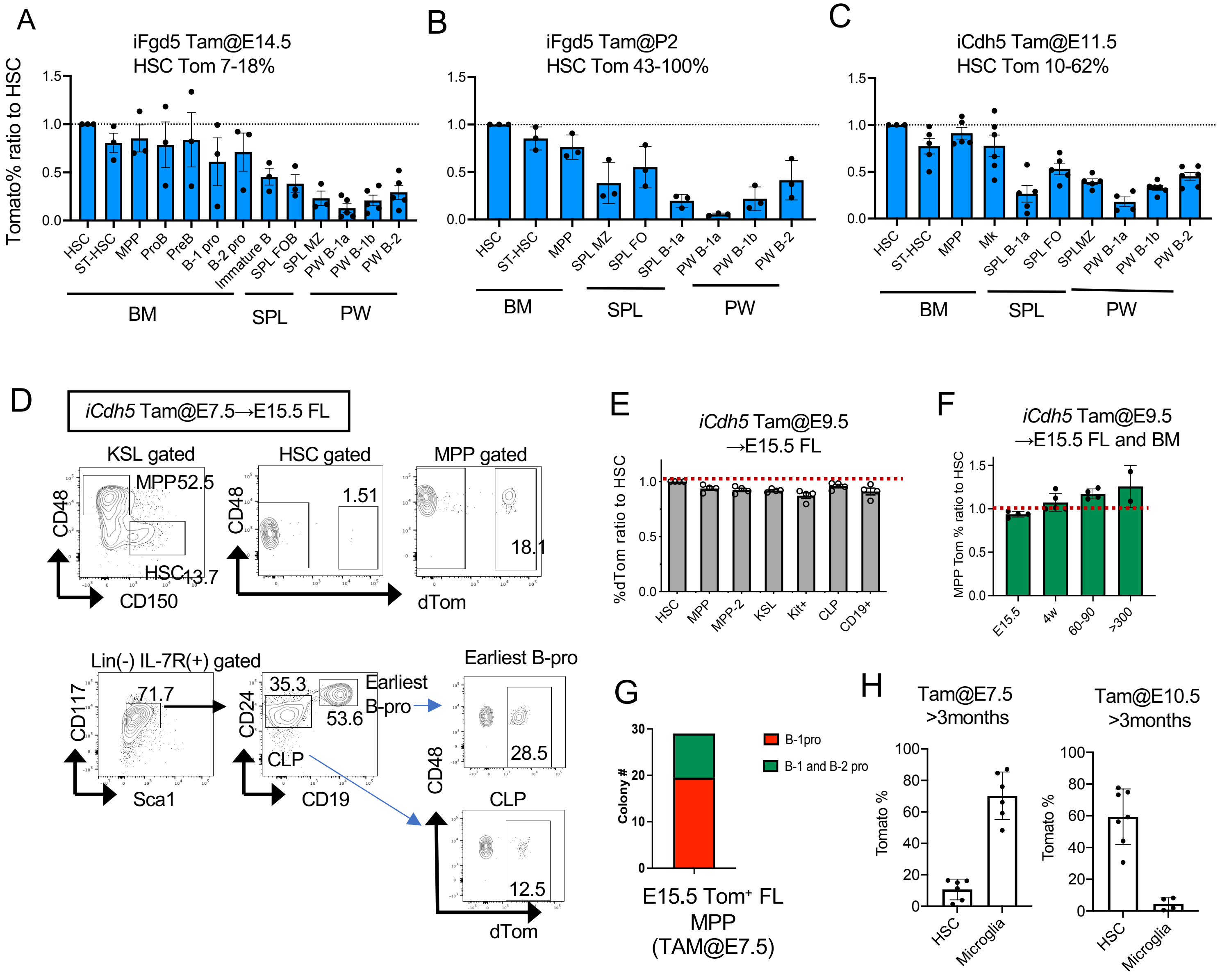


**Extended Data Fig. 1 HSC- and EC-lineage studies revealed HSC-independent definitive hematopoiesis persist into adult life**

The relative Tomato% ratios of HSPCs and B cell subsets to LT-HSCs at >300 days after birth when TAM was injected at E14.5 (A) or P2 (B) into iFgd5 mice, or at E11.5 into iCdh5 mice (C). N=3-7 for each TAM injection. BM: bone marrow, SPL: spleen, PW: peritoneal wash. (D) Representative FACS plots of LT-HSC, MPP, CLP and the earliest B-progenitors in E15.5 FL of *iCdh5* embryos injected with TAM at E7.5 (n=6). (E) The relative Tomato% ratio of each HSPC to LT-HSCs in E15.5 FL of iCdh5 embryos when TAM was injected E9.5 (n=4). (F) The relative Tomato% ratio of MPPs to LT-HSCs in the E15.5 FL and post-natal BM at different time points (4w, days 60-90, days 300<, n=3-4 at each time point). (G) B-cell progenitor colony forming cell counts from 500 Tom$^{+}$ MPPs from E15.5 FL of *iCdh5* mice that was injected with TAM at E7.5 (n=3). (H) Tomato% of HSCs and brain macrophates when Tam was injected into iCdh5 mice at E7.5 and E10.5 (n=7).

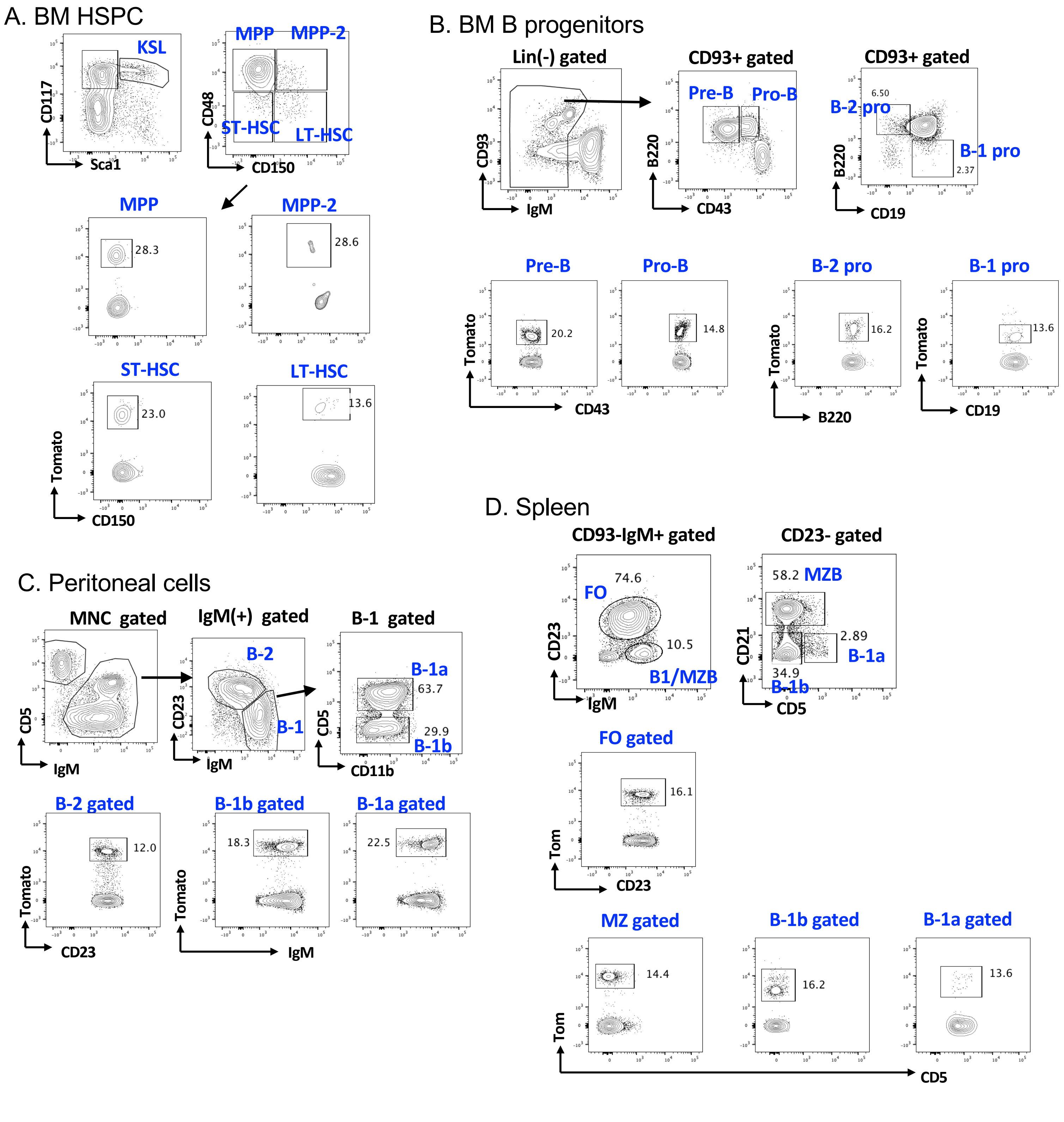


**Extended Data Fig. 2 Gating strategies of HSPC and B-lymphoid subsets**
Representative FACS plots for BM HSPC (A), BM B-progenitors (B), peritoneal B cell subsets (C), and spleen B cell subsets (D). The data is from *iCdh5* mice injected TAM at E8.5 and analyzed 6 months after birth.

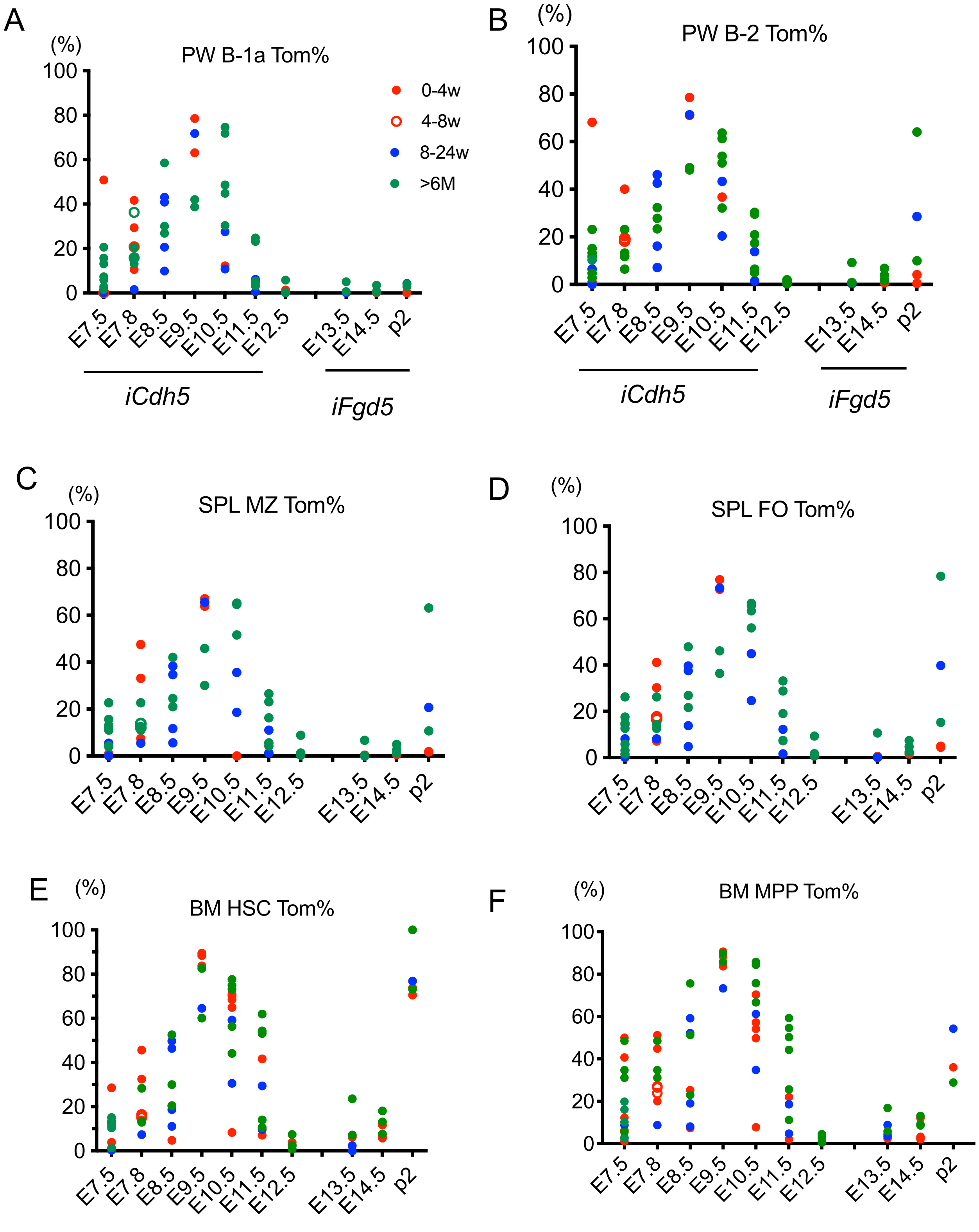


**Extended Data Fig. 3. Tomato % in B-cell subsets and HSPCs after TAM injection into *iCdh5* and *iFgd5* mice at different time points.**

TAM was injected into iCdh5 or iFgd5 mice at different time points and mice were analyzed at either 0-4 weeks (red circle), 4-8 weeks (open red circle), 8-24 weeks (blue circle), and more than 6 months (green circle). PW: peritoneal wash, SPL: spleen, BM: bone marrow, MZ: mariginal zone B cell, FO: follicular B cell, HSC: hematopoietic stem cell, MPP: multipotent progenitor

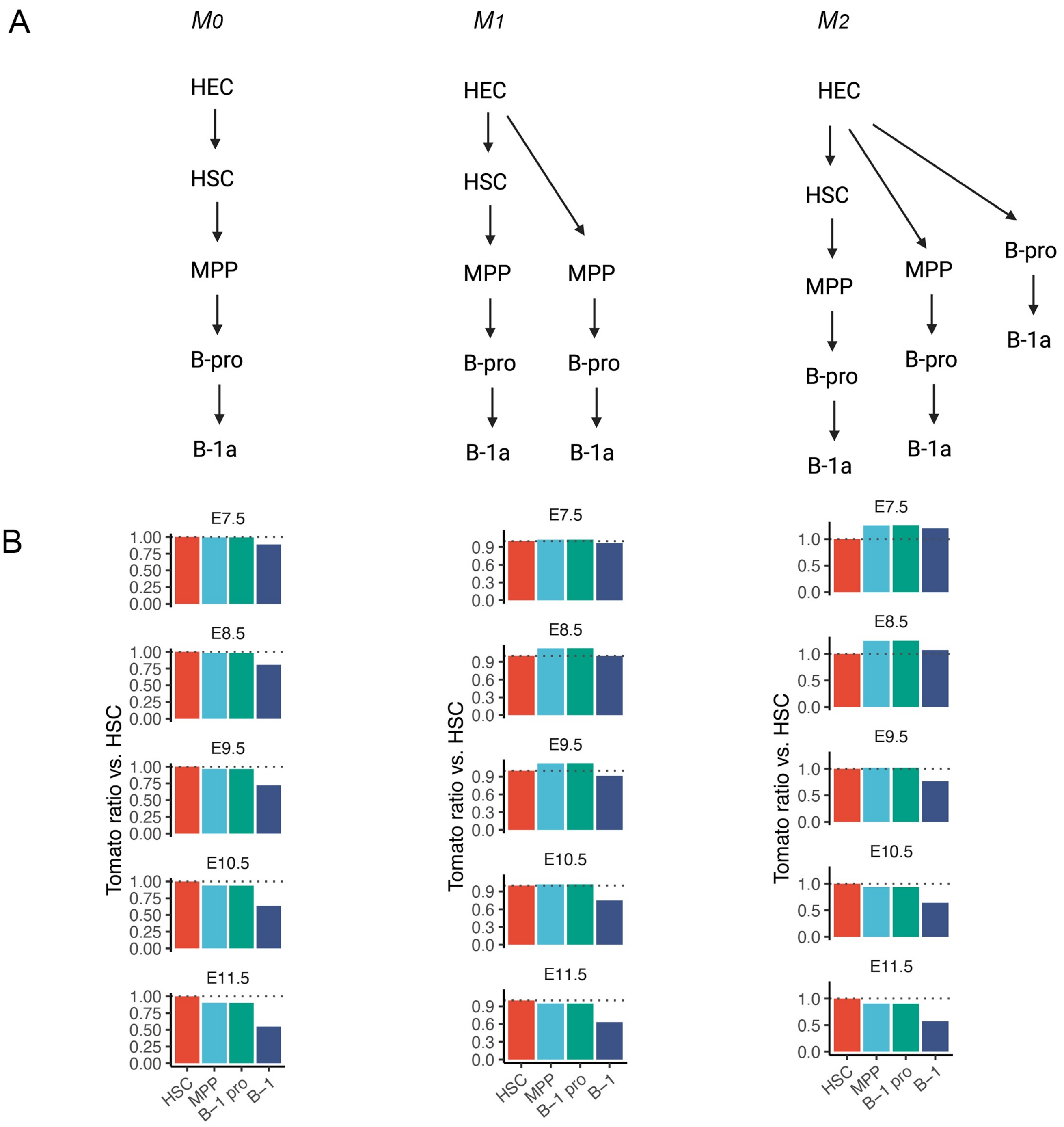


**Extended Data Fig. 4. Simulated cell differentiation labeling ratios under competing lineage tree models.**

Cell differentiation was simulated using a mathematical model (see Methods) in order to reproduce the label tracing results from the iCdh5 mouse (Fig. 1I-K). The model was initialized with a population of Tomato- hemogenic ECs, and these cells became Tomato+
due to tamoxifen injection at the indicated embryonic days. Subsequently, Tomato labels were propagated to downstream progeny cell types according to the indicated lineage tree models. (A) Lineage tree models $M_0$, $M_1$, and $M_2$. (B) Tomato ratios relative to HSC for each cell type and timing of tamoxifen injection, under the indicated lineage tree model.

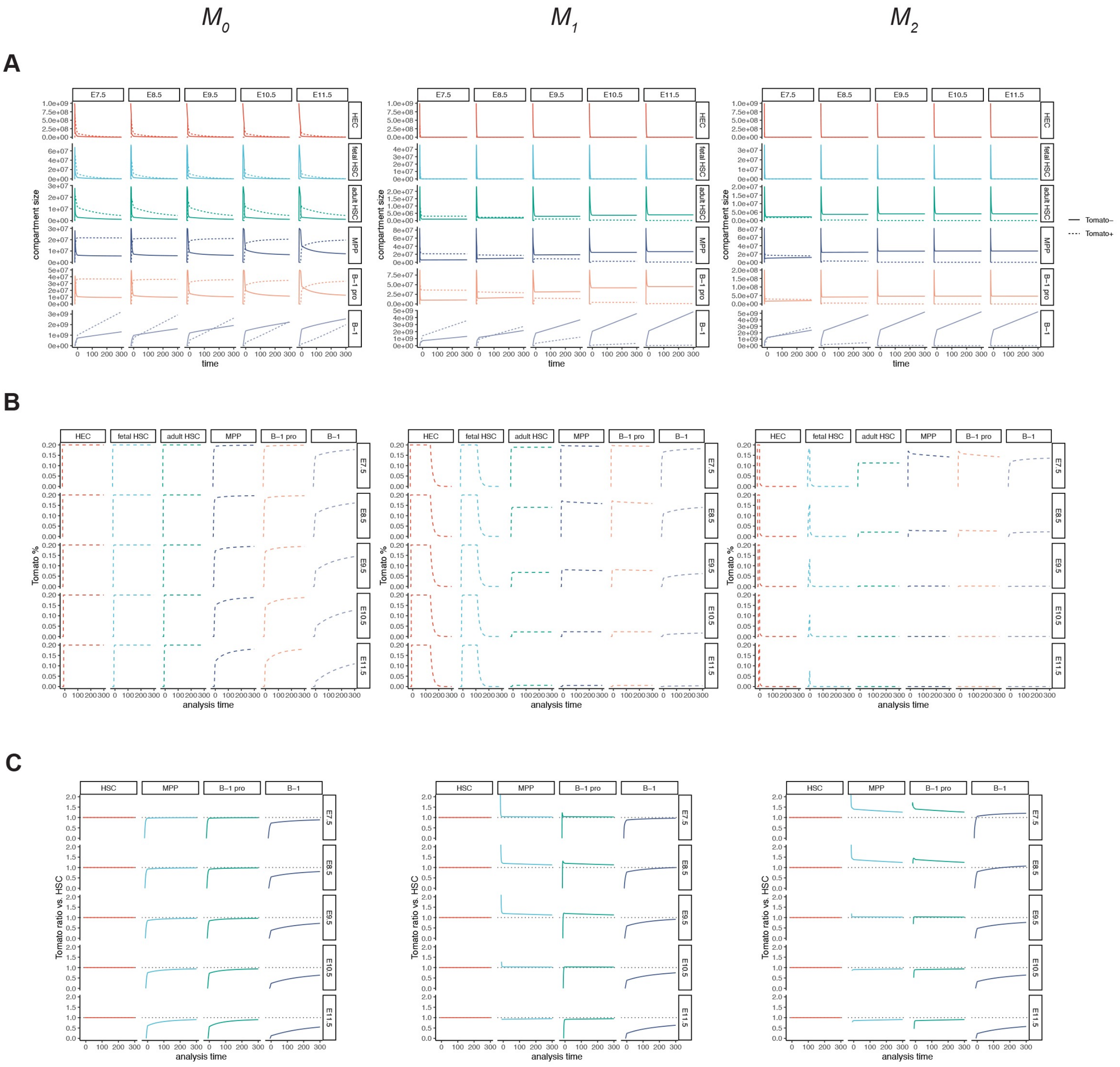


**Extended Data Fig. 5. Simulated cell differentiation trajectories under competing lineage tree models.**

Cell differentiation was simulated using a mathematical model (see Methods) in order to reproduce the label tracing results from the iCdh5 mouse (Fig. 1I-K). The model was initialized with a population of Tomato- hemogenic ECs, and these cells became Tomato+ due to tamoxifen injection at the indicated embryonic days. Subsequently, Tomato labels were propagated to downstream progeny cell types according to lineage tree model M0, M1, or M2 as shown in Extended Data Fig. 4.

The simulated trajectories of (A) compartment sizes, (B) percentage of Tomato+ cells, and (C)Tomato ratio relative to HSC are plotted across time after birth for each cell type and tamoxifen injection timing. The HSC compartment was simulated as two distinct latent subpopulations (fetal and adult HSC), but they were combined as one compartment for the Tomato ratio calculation.

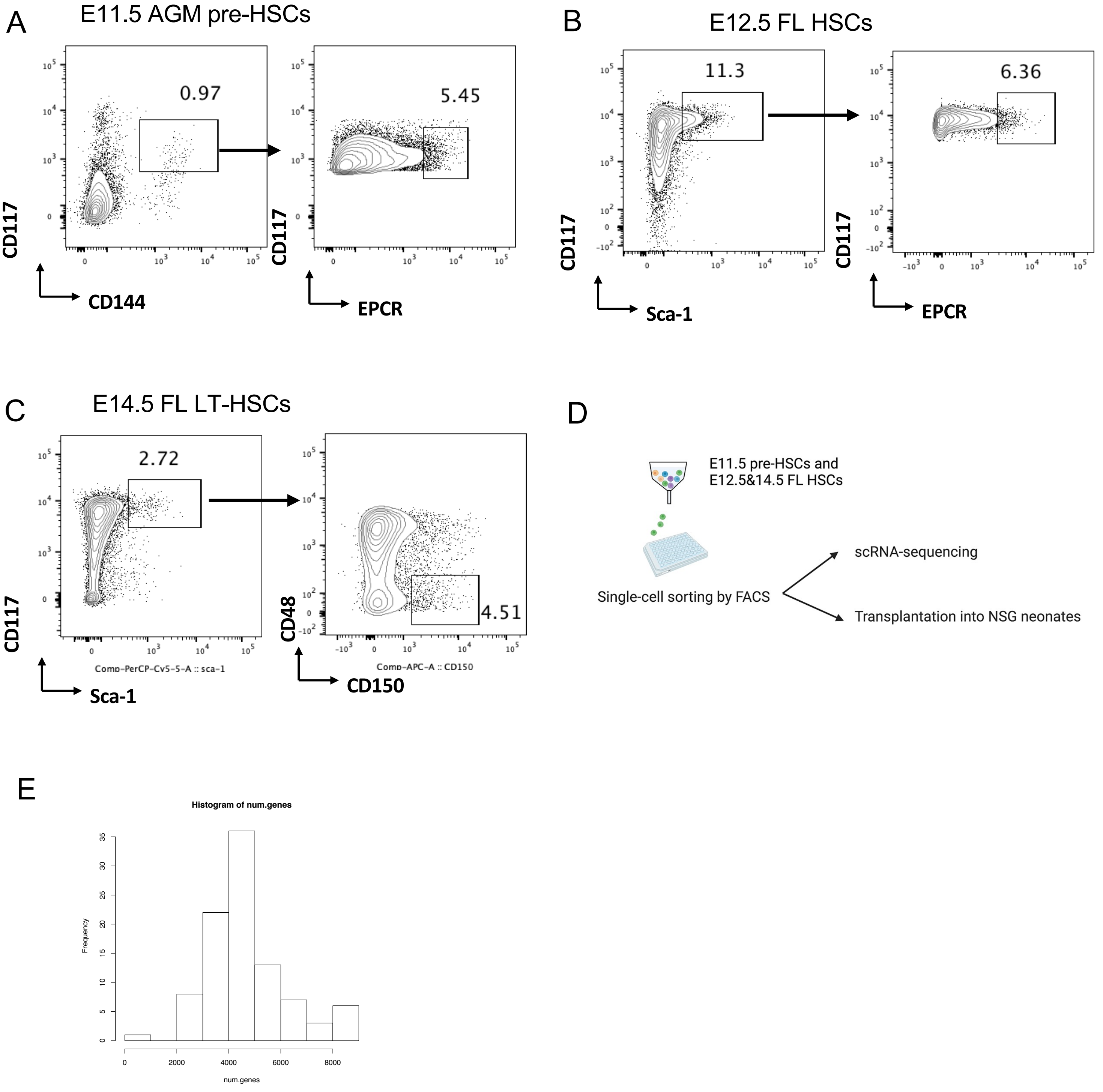


**Extended Data Fig. 6. Gating strategies of pre-HSC and FL HSCs at different stages.** Representative FACS plots for E11.5 pre-HSCs (A), E12.5 FL HSCs(B), and E14.5 FL HSCs. (D) The experimental design for scRNA-seq. While 96 single cells were sorted, 5-10 pre-HSCs or HSCs were transplanted into NSG mice to validate the biological functions. (E) Histogram of numbers of genes expressed in each cell (read count >10). We filter the cells which have less than 2000 genes (read counts >10) and genes (read counts >10) which are detected in less than 10 cells. At last, 95 cells and 11,814 genes are used for further analysis.

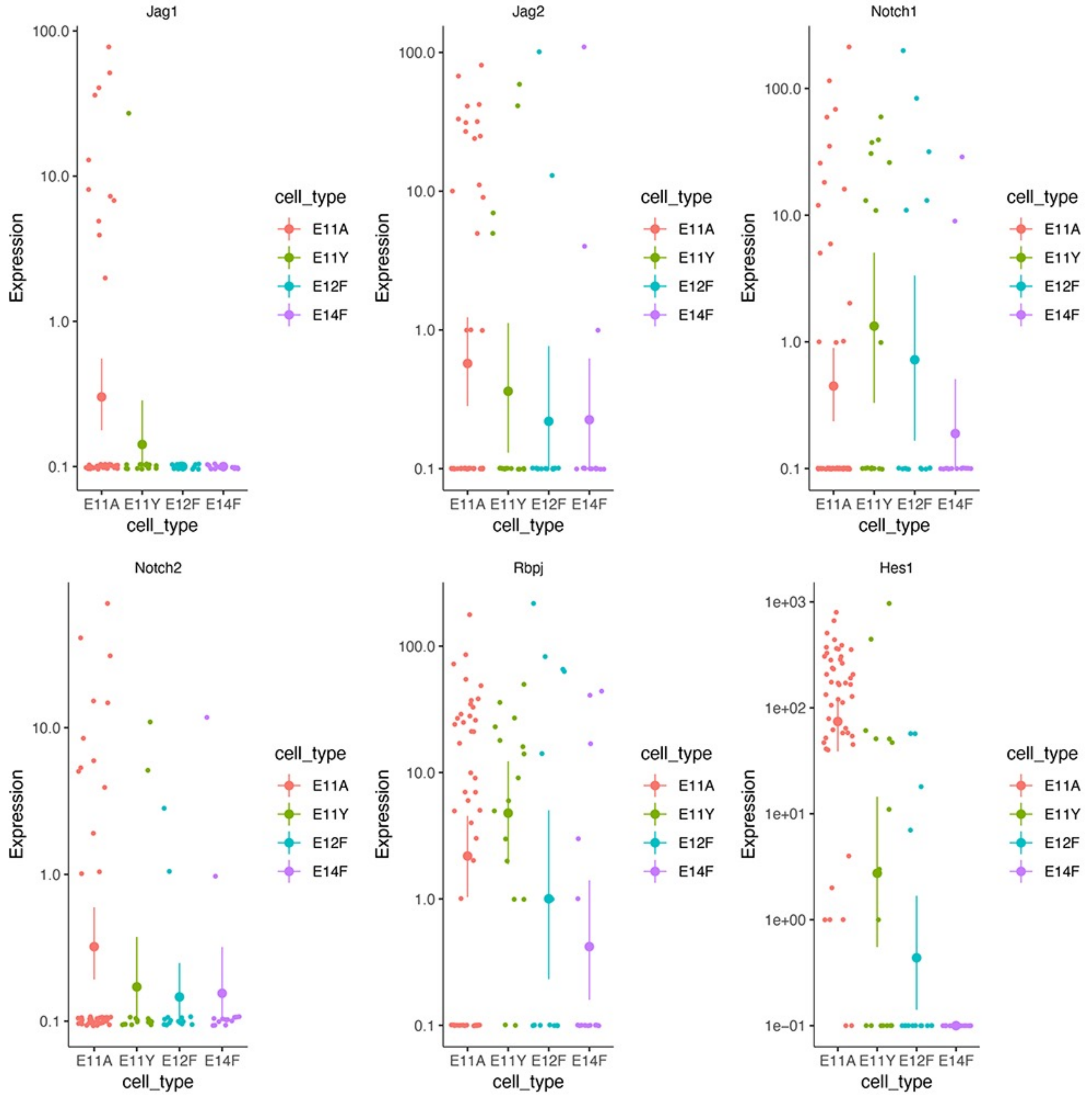


**Extended Data Fig. 7. Notch signaling is reduced during the transition from pre-HSCs to HSCs.**

Expressions of Notch related genes are shown in each single cells from VE-cadherin$^{+}$c-kit$^{+}$CD45$^{+}$ EPCR$^{+}$ cells from E11.5 AGM and YS, Ter119-CD45$^{+}$c-kit$^{+}$Sca-1$^{+}$EPCR$^{+}$E12.5 FL HSCs, and lin$^{-}$Sca-1$^{+}$c-kit$^{+}$ CD48$^{-}$CD150$^{+}$E14.5 FL HSCs.

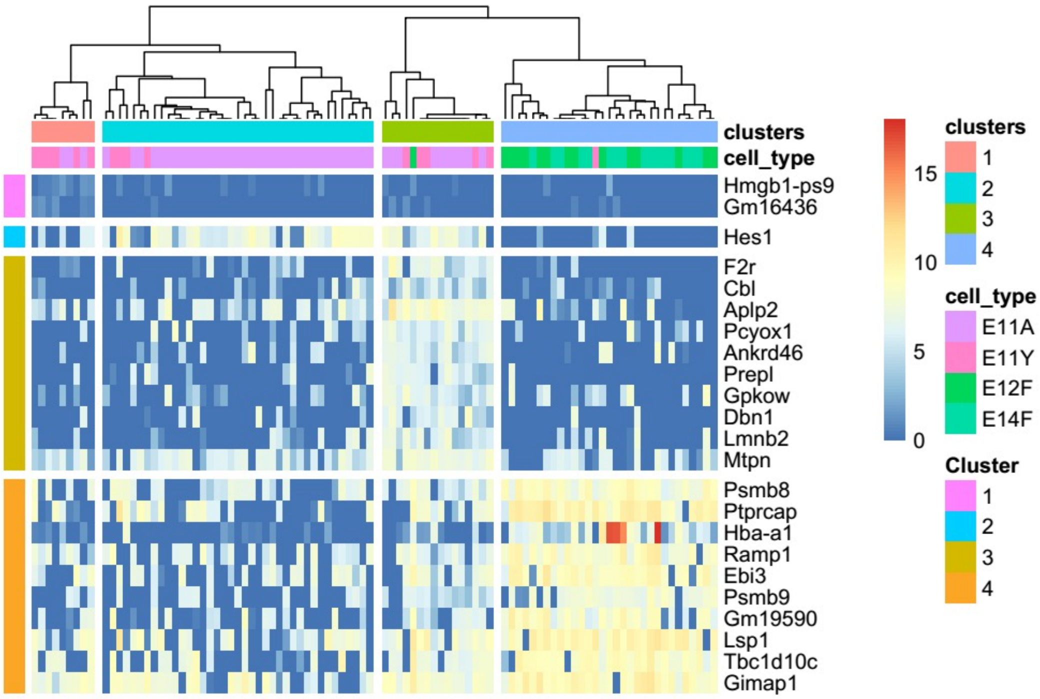


**Extended Data Fig. 8.** Heatmap of normalized scRNA-seq expression of markers (the area under the ROC curve (auroc) > 0.85 and the adjusted p-values <0.01). The top 10 differentially expressed genes are depicted

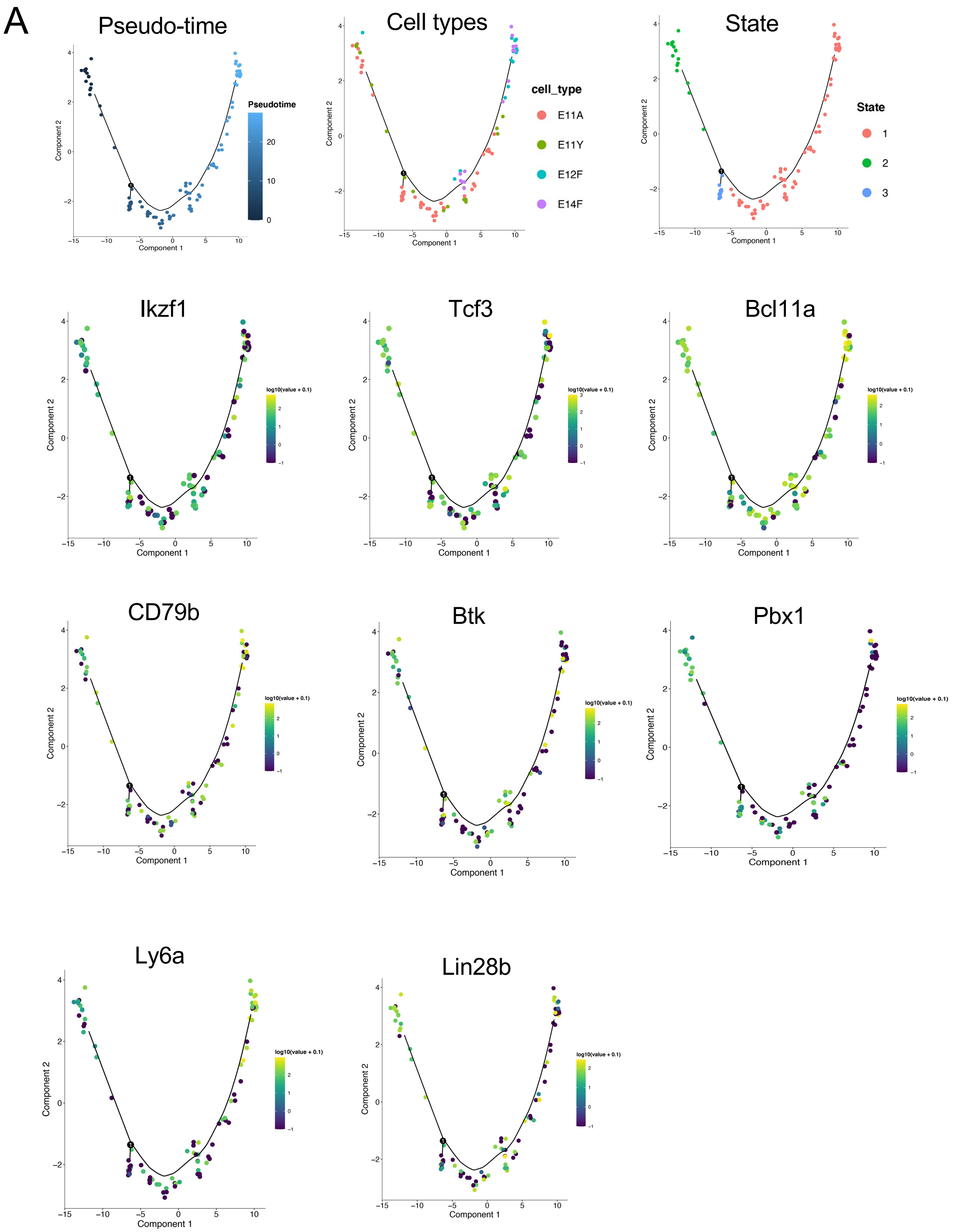


**Extended Data Fig. 9. Gene expressions in each pre-HSC and FL HSC.**
Pseudo-time-ordering trajectory of scRNA-seq data and each cell was colored by cell type (middle) and states (right) using Monocle2. Some HSC-related and early B-cell commitment related genes are depicted.
